# Is understanding the structure-function of microbial community of sludge suitable using DNA-based molecular methods?

**DOI:** 10.1101/536508

**Authors:** Ping Wang, Zhen Wang, Jing Ma

**Affiliations:** School of Chemistry and Chemical Engneering, Zhoukou Normal University, Zhoukou 466001, China; Yellow River Institute of Hydraulic Research, Zhengzhou 450003, China; Key Laboratory of Yellow River Sediment Research, MWR, Zhengzhou 450003, China

**Keywords:** running sludge, idle sludge, total microbial community, active microbial community, Illumina MiSeq

## Abstract

Analysis the similarities between total and active microbial community is crucial to evaluate whether understanding the structure-function of microbial community of different types sludge is suitable using DNA-based molecular methods. In this study, procaryotic communities in different types sludge samples (including primary sludge, excess sludge, mixed sludge, and digested sludge) were evaluated using DNA-based and RNA-based Illumina MiSeq. Results showed that the similarities between total and active procaryotic communities of all different types sludge were considerable high. Meanwhile, the similarity in running sludge higher than that in idle sludge, because of the Jarccard’s coefficient of digested sludge was the highest in all types sludge samples. This study indicates that the DNA-based molecular biology technology could be applicable to profile procaryotic community structure and function in sludge samples, and this study adds the understanding of total and active procaryotic communities of sludge samples.

## 1. Introduction

Activated sludge technology is currently the most broadly implemented biological method for biomass conversion in wastewater treatment plants (WWTPs) (Kim et al., 2003; Ye and Zhang, 2013; Korzeniewska and Harnisz, 2018). However, this process produced vast quantities of highly organic waste activated sludge, and this by-product mass continues to increase with the expansion of population and industry (Cui and Jahng, 2006; Zhang et al., 2010). Sludge disposal by landfill or incineration may no longer be feasible in the near future due to land scarcity, high waste charge and increasing stringent environmental control regulations (Weemaes et al., 2000; Wong et al., 2013; Wang et al., 2018). Therefore, the strategy for sludge management is shifting towards its reutilization as a potential source for renewable energy (Mossakowska et al., 1998; Rulkens, 2007). In this regard, the anaerobic digestion while producing renewable energy in the form of biogas (Riviere et al., 2009; Sundberg et al., 2013).

The processes of wastewater treatment using activated sludge technology and sludge disposal using anaerobic digestion depend on the synergistic interactions of microorganisms (Vanwonterghem et al., 2014; Wagner et al., 2002; Zakrzewski et al., 2012; Zhang et al., 2012). The culturing methods have been a very direct and effective way to characterize the microbial community (Neilson, 1978). However, most of the microorganism in complex environment cannot be cultured in an artificial medium in the laboratory and the diversity of the uncultured microorganism is quite considerable (Riviere et al., 2009). The molecular methods, such as polymerase chain reaction (PCR)-denaturing gradient gel electrophoresis (Supaphol et al., 2011; Ye and Zhang, 2011), terminal restriction fragment length polymorphism (Liu et al., 1997), cloning (Rincón et al., 2008; Schuppler et al., 1995), fluorescence *in situ* hybridization (Braguglia et al., 2012; Erhart et al., 1997) etc., greatly promoted our understanding of the microbial community. Typically, these molecular methods are gained through directly extracted DNA, but in the case of sludge samples, the problem is that this DNA contains not only the active microbial populations of interest but also genes of dormant, dead, and degraded cells. Because of these non-target pools that archive past and inactive genetic diversity, sludge DNA has been claimed to represent sludge history. RNA, on the other hand, is more responsive with a higher turnover rate within cells. RNA-based molecular methods can profile the structure-function links of microbial communities of environmental samples, which is a central theme of microbial ecology since its beginning (Brettar et al., 2012b; Sudhagar et al., 2018). However, RNA-based molecular methods are difficult to operate due to the easy degradability of RNA. If the two nucleic acid pools are extracted from the same environmental sample and analyzed with the same methodology, the structures of the DNA-based (likely including partly past) and RNA-based (likely currently viable and potentially more active) communities can be compared, and then evaluate the similarities between total and active microbial community. PCR-based community Illumina sequencing such as MiSeq is one popular high throughput sequencing systems due to its high quality and low cost. MiSeq of DNA-based and RNA-based has been successfully used in investigating total and active microbial community in various samples, such as forest soil (Baldrian et al., 2012a), land farming soil (Mikkonen et al., 2014), sediment from hydrothermal vent field (Lanz et al., 2011), and seawater (Brettar et al., 2012a). There are also several studies applying these methods in exploring total and active microbial community in anaerobic digestion reactors (Zakrzewski et al., 2012; Franchi et al., 2018). However, analysis of the similarities between total and active microbial community in different types sludge using Illumina MiSeq is still very limited.

The purpose of this study was to evaluate that whether understanding the structure-function of microbial community of different types sludge using DNA-based molecular methods is suitable, according to analysis the similarities between total and active microbial community using DNA-based and RNA-based Illumina MiSeq. Procaryotic communities in different types sludge samples (including primary sludge, excess sludge, mixed sludge, and digested sludge) were evaluated in this study. For each sludge sample, we extracted both DNA and RNA, and then prepared and pyrosequenced PCR amplicons from both extractions.

## 2. Materials and methods

### 2.1 Sludge digestion system description and sampling

In this study, all sludge samples were taken from a full-scale WWTP and its subsequent sludge anaerobic digestion system. Sampling was performed on May 28, and got 8 sludge samples. These samples were respectively primary sludge (PS) from primary settling tank, excess sludge (ES) from secondary settling tank, primary digested sludge I (PDS-I) from primary digester I, primary digested sludge II (PDS-II) from primary digester II, primary digested sludge III (PDS-III) from primary digester III, secondary digested sludge (SDS) from secondary digester, concentrated sludge (CS) (PS and ES mixed), and dewatering sludge (DS) (SDS and PS mixed, and then dewatered). All samples were collected in triplicated and were keep immediately in liquid nitrogen before being transported to the laboratory. Samples were stored at −80°C until DNA and RNA extraction.

### 2.2 Nucleic acid extraction and reverse transcription

DNA was extracted from 2 mL sludge (except DS, 0.5 g) using a PowerSoil DNA isolation kit (MoBio Laboratories, Carlsbad, CA, USA) according to the manufacturer’s instructions. DNA was diluted 0-, 5-, and 10-fold for PCR, to adjust the DNA concentration to the proper condition for PCR, and to reduce the influence of the PCR inhibitors (e.g., humic substances).

RNA extraction was performed as previously (Wang et al., 2011). RNA was extracted from 2.0 g sludge using RNA PowerSoil total RNA isolation Kit (MoBio, Carlsbad, CA, USA). Extracted RNA was purified using Illustra MicroSpin Columns S-400 HR (GE Healthcare, Buckinghamshire, UK) to remove humic substances, and the remaining DNA was removed using a Turbo DNA-free kit (Applied Biosystems, Foster City, CA, USA). RNA was further purified and concentrated using an RNA Clean & Concentrator-5 kit (Zymo Research, Orange, CA, USA). The amount of purified RNA was measured using a NanoDrop 1000 (NanoDrop Products, Wilmington, DE, USA). The absence of DNA carryovers in the RNA samples was verified by PCR targeting the 16S rRNA gene without reverse transcription. Purified RNA was reverse transcribed using random hexamers and PrimeScript Reverse Transcriptase (Takara Bio, Otsu, Shiga, Japan) with 20 U of SUPERase-In RNase Inhibitor (Applied Biosystems). Synthesized cDNA was diluted 10-fold for PCR. Replicate DNA and cDNA samples were pooled and used for Illumina MiSeq Sequencing.

### 2.3 Barcoge amplicon MiSeq sequenving

The V4 region of 16S rRNA gene was targeted and amplified for all cDNA and DNA libraries prepared using the universal forward primer 515F (5’-barcode-GTGCCAGCMGCCGCGGTAA-3’) and the universal reverse primer 909R (5’-CCCCGYCAATTCMTTTRAGT −3’). The barcode is a sixteen-base sequence unique to each sample. PCR reactions were performed in triplicate in a 50-μL mixture containing 5 μL 10×PCR buffer, 2.0 μL 2.5 mM MgCl_2_, 2.0 μL 2.5 mM dNTP, 2.0 μL 10 mM forward primer, 2.0 μL 10 mM reverse primer, 0.5 μL TransStart Fast Pfu DNA Polymerase (TianGen, China), 10 ng DNA (or cDNA) and ddH_2_O. The PCR amplification program included initial denaturation at 94°C for 3 min, followed by 30 cycles of 94°C for 40 s, 56°C for 60 s, and 72°C for 60 s, and a final extension at 72°C for 10 min. Then, the PCR products of 3 replicated were mixed and purified with the EZNA® Cycle-Pure Kit (Omega Bio-tek Inc, Doraville, GA, USA) and quantified using NanodropND-1000 UV-Vis Spectrophotometer (NanoDrop Technologies, Wilmington, DE, USA). All samples were pooled together with equal molar amount from each sample. The sequencing samples were prepared using TruSeq DNA kit according to manufacture’s instruction. The purified library was diluted, denatured, re-diluted, mixed with PhiX (equal to 30% of final DNA amount) as described in the Illumina library preparation protocols, and then applied to an Illumina Miseq system for sequencing with the Reagent Kit v2 2×250 bp as described in the manufacture manual.

### 2.4 Processing and analyzing of sequencing data

Raw FASTQ files were de-multiplexed and quality-filtered using QIIME. The sequence data were processed using QIIME Pipeline–Version 1.7.0 (http://qiime.org/tutorials/tutorial.html) with the following criteria: (i) The 300-bp reads were truncated at any site that obtained an average quality score of <20 over a 10-bp sliding window, and the truncated reads shorter than 50 bp were discarded; (ii) exact barcode matching, two nucleotide mismatch in primer matching, and reads containing ambiguous characters were removed; and (iii) only overlapping sequences longer than 10 bp were assembled according to their overlapped sequence. Reads that could not be assembled were discarded. Sequences were clustered into operational taxonomic units (OTUs) at a 97% identity threshold. The aligned 16S rRNA gene sequences were used for chimera check using the Uchime algorithm (Edgar et al., 2011).

The rarefaction analysis based on Mothur v.1.21.1 (Schloss et al., 2009) in QIIME was conducted to reveal the diversity indices, including the Chao1, coverage, and Shannon diversity indices. The beta diversity analysis was performed using UniFrac (Lozupone et al., 2011) to compare the results of the principal component analysis (PCA) and cluster analysis (CA) using the Vengan and Permute package in R (x64 3.2.2). Jarccard’s coefficient was used to indicate the similarity between active and total microbial communities in same sludge.

## 3. Results

### 3.1 Sequencing results and diversity indices

As shown in Table 1, a total of 176216 (DNA-based) and 155563 (RNA-based) effective sequences were obtained from eight samples through Illumina MiSeq analysis. In order to do the comparison at the same sequencing depth, 9295 effective sequences were extracted from each sample to do diversity analysis. The phylogenetic OTUs of each library derived DNA ranged from 553 to 1958, and which derived RNA ranged from 313 to 937. The diversity of RNA-derived communities was less than that of the DNA-derived communities, particularly in CS and DS (Table 1). Among them, OTUs numbers of CS and DS derived RNA in proportion of these derived RNA were 37% and 23%, respectively. However, the value was up to 77% in PDS. OTUs numbers of all sludge samples showed that the presence of a large variation in the total number OTUs in different types sludge samples. The OTUs densities derived DNA of all samples showed that OTUs number of PDS was the lowest and OTUs number of CS was the highest. The OTUs densities derived RNA of all samples showed that OTUs number of DS was the lowest and OTUs number of ES was the highest. These results showed that active microbial community in the proportion of total microbial community are very different in different types sludge samples. Moreover, the calculation of the Chao1, coverage, and Shannon index derived DNA confirmed that CS has the highest diversity and PDS has the lowest diversity in all sludge samples. However, these indexes derived RNA confirmed that ES has the highest diversity and DS has the lowest diversity in all sludge samples. All coverage indexes indicated that most communities were accounted for employing Illumina MiSeq.

**Table 1.** Analysis of the prokaryotic diversity indices and richness estimators

| Sample | <sup>a</sup>Sequences | <sup>b</sup>Sequences | OTUs | Shannon<br>(nat) | Chao1 | Coverage<br>(%) |
| --- | --- | --- | --- | --- | --- | --- |
| PS-D | 9295 | 9295 | 1398 | 4.26 | 3214 | 82.02 |
| PS-R | 10758 | 9295 | 772 | 2.94 | 1775 | 95.48 |
| ES-D | 14457 | 9295 | 1754 | 4.63 | 3702 | 77.74 |
| ES-R | 16238 | 9295 | 937 | 3.15 | 1947 | 94.26 |
| CS-D | 21676 | 9295 | 1958 | 4.79 | 3917 | 80.24 |
| CS-R | 23682 | 9295 | 720 | 2.71 | 1568 | 90.85 |
| PDS-I-D | 26466 | 9295 | 553 | 2.59 | 1418 | 92.73 |
| PDS-I-R | 17683 | 9295 | 421 | 2.44 | 1069 | 95.21 |
| PDS-II-D | 29473 | 9295 | 649 | 2.68 | 1724 | 91.17 |
| PDS-II-R | 27112 | 9295 | 478 | 2.51 | 1193 | 93.43 |
| PDS-III-D | 32488 | 9295 | 594 | 2.62 | 1572 | 93.27 |
| PDS-III-R | 15897 | 9295 | 452 | 2.48 | 1123 | 94.96 |
| SDS-D | 23265 | 9295 | 673 | 2.73 | 1795 | 90.83 |
| SDS-R | 20372 | 9295 | 431 | 2.46 | 1086 | 95.06 |
| DS-D | 19096 | 9295 | 1339 | 4.67 | 3692 | 85.04 |
| DS-R | 23821 | 9295 | 313 | 2.05 | 983 | 94.43 |

<sup>a</sup>Sequences: effective sequence of each sample;
<sup>b</sup>Sequences: in order to do the comparison at the same sequencing depth, 9295 effective sequences were extracted from each sample to do diversity analysis;
Abbreviations: PS, primary sludge; ES, excess sludge; CS, concentrated sludge; PDS-I, primary digested sludge I; PDS-II, primary digested sludge II; PDS-III, primary digested sludge III; SDS, secondary digested sludge; and DS, dewatering sludge; -D, DNA-based; -R, RNA-based.

Results of CA (as shown in Figure 1) and PCA (as shown in Figure 2) showed that microbial community derived RNA and DNA of the same sludge sample are very similar, and aerobic environment samples (such as PS, ES, and CS) with anaerobic environment sample (e.g., PDS and SDS) were in different partitions. For example, PDS and SDS were clustered together due to all of them in anaerobic environment.

**Figure 1.**
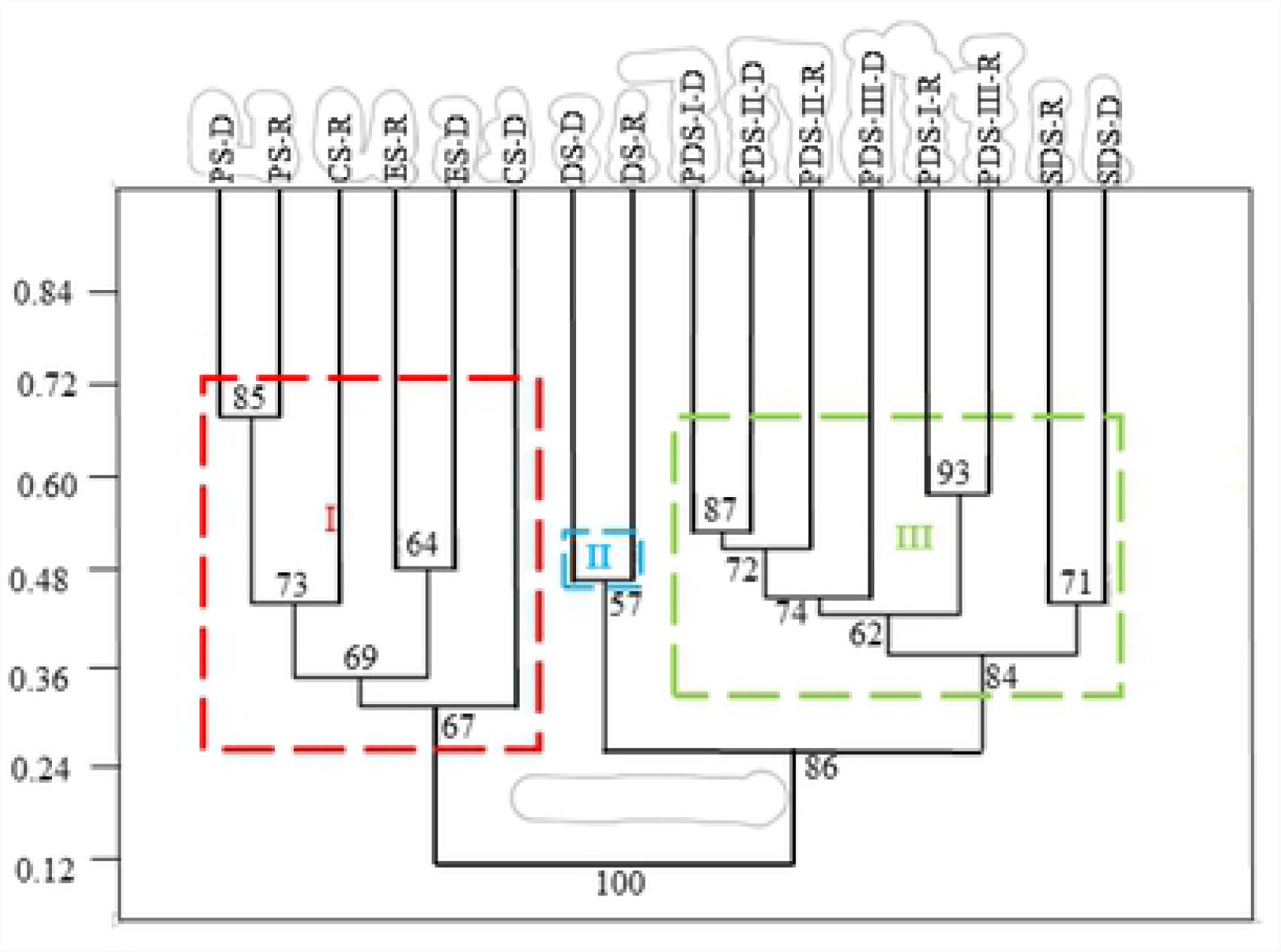
Clustering analysis of the similarities among samplcs’s prokaryotic communities using Weighted-UniFrac. Abbreviations: PS, primary sludge; ES, excess sludge; CS, concentrated sludge; PDS-I, primary digested sludge I; PDS II, primary digested sludge II; PDS-III, primruy digested sludge Ill; SDS, secondary digested sludge; and DS, dcwatering sludge; −D, DNA based; −R, RNA-based.

### 3.2 Prokaryotic community compositions

We applied Illumina Miseq to evaluate whether and how DNA-based (16S rRNA gene) and RNA-based (16S rRNA gene) microbial community derived from the same sludge sample differed, to evaluate differences and similarities in microbial community (DNA-based or RNA-based) derived from different sludge samples. As shown in Table 2, microbial communities derived RNA and DNA in sludge samples were predominated by bacteria, with abundances between 80.80% and 92.00%, while abundances of archaea were between 0.20% and 4.70%. The ratio of archaeal sequences was higher in digested sludge samples (mean 4.13% and 2.93% derived DNA and RNA) than in other sludge samples.

#### 3.2.1 Prokaryotic community composition in PS

Total prokaryotic community and active prokaryotic community in PS were clustered together with high similarity (Jarccard’s coefficient, 81.78%) at phylum level. As shown in Figure 3, *Firmicutes* (36.71% and 38.92%), *Proteobacteria* (35.20% and 40.52%), *Bacteroidetes* (10.58% and 8.96%), *Actinobacteria* (9.64% and 6.43%), *Euryarchaeota* (3.38% and 0.94%), and *Chloroflexi* (1.03% and 0.83%) were dominated in total prokaryotic community and active prokaryotic community. Among them, *Firmicutes* and *Proteobacteria* were highly abundant in two prokaryotic communities. Meanwhile, abundances of those 2 phyla in active prokaryotic community were higher than these in total prokaryotic community. However, abundances of other phyla in active prokaryotic community were lower than these in total prokaryotic community.

At genus level, total prokaryotic community and active prokaryotic community in PS were also similar (Jarccard’s coefficient, 63.54%). As shown in Figure 4, *Trichococcus* (18.51%), *Arcobacter* (8.83%) and *Comamonadaceae*(family) (means ‘an unknown genera belonged to the family *Comamonadaceae*’, the same below) (7.28%) were the top 3 genera in total prokaryotic community, and *Trichococcus* (22.72%), *Arcobacter* (9.31%), and *Acinetobacter* (8.41%) were the top 3 genera in active prokaryotic community.

#### 3.2.2 Prokaryotic community composition in ES

Total prokaryotic community and active prokaryotic community in ES were clustered together with high similarity (Jarccard’s coefficient, 82.96%) at phylum level. As shown in Figure 3, *Actinobacteria* (26.26% and 28.39%), *Proteobacteria* (27.51% and 22.64%), *Chloroflexi* (21.95% and 22.45%), *Bacteroidetes* (12.68% and 9.54%), *Firmicutes* (4.37% and 3.51%), *Planctomycetes* (2.84% and 1.04%), TM7 (1.74% and 3.44%), and *Chlorobi* (0.79% and 1.10%) were dominated in total prokaryotic community and active prokaryotic community. Among them, *Actinobacteria, Proteobacteria*, and *Chloroflexi* were highly abundant in two prokaryotic communities. Meanwhile, abundances of those 3 phyla in active prokaryotic community were higher than these in total prokaryotic community. In addition to 3 phyla, TM7 and *Chlorobi* in active prokaryotic community were a little higher than these in total prokaryotic community. However, abundances of other phyla in active prokaryotic community were lower than these in total prokaryotic community.

At genus level, total prokaryotic community and active prokaryotic community in PS were also similar (Jarccard’s coefficient, 67.87%). As shown in Figure 4, DRC31(order) (10.01%), *Actinomycetales*(order) (7.16%) and *Microthrixaceae*(family) (7.03%) were the top 3 genera in total prokaryotic community, and DRC31(order) (9.64%), *Microthrixaceae*(family) (7.41%), and *Candidatus M. parvicella* (6.78%) were the top 3 genera in active prokaryotic community.

#### 3.2.3 Prokaryotic community composition in digested sludge

Total prokaryotic community and active prokaryotic community in digested sludge were clustered together with high similarity (Jarccard’s coefficient, 81.81-85.68%) at phylum level. As shown in Figure 3, our results showed that prokaryotic communities were very similar in three parallel anaerobic digesters, suggesting that experimental results in this study are repeatable and credible. Moreover, prokaryotic community in secondary digester (normal temperature digestion) was similar to these in primary digesters (mesophilic digestion). *Verrucomicrobia* (40.43-45.72%), WWE1 (10.46-17.24%), *Chloroflexi* (5.60-11.86%), *Bacteroidetes* (7.35-13.22%), *Firmicutes* (4.96-9.67%), *Proteobacteria* (2.27-8.34%), TM7 (0.78-2.11%), *Euryarchaeota* (0.96-4.36%), OD1 (1.50-1.94%), and *Synergistetes* (1.31-3.23%) were dominated in total prokaryotic community and active prokaryotic community in digested sludge. Among them, abundances of *Verrucomicrobia, Proteobacteria, Euryarchaeota*, OD1, and *Synergistetes* in active prokaryotic community were higher than these in total prokaryotic community. However, abundances of other phyla in active prokaryotic community were lower than these in total prokaryotic community. Moreover, *Verrucomicrobia*, WWE1, *Chloroflexi, Bacteroidetes*, and *Firmicutes* were highly abundant in two prokaryotic communities.

At genus level, total prokaryotic community and active prokaryotic community in PS were also similar (Jarccard’s coefficient, 73.82-77.49%). As shown in Figure 4, LD1-PB3(order) (30.43-33.72%), W22 (6.85-10.80%), and T78 (6.21-10.49%) were the top 3 genera in total prokaryotic community, and those also were the top 3 phyla in active prokaryotic community.

#### 3.2.4 Prokaryotic community composition in mixed sludge

At phylum level, the similarities between total prokaryotic community and active prokaryotic community in mixed sludge were lower than that in single sludge. Jarccard’s coefficient of CS and DS were respectively 74.41% and 72.10%. Just like at phylum level, the similarities between total prokaryotic community and active prokaryotic community in mixed sludge were lower than that in single sludge because that Jarccard’s coefficient of CS and DS were respectively 44.37% and 41.72%.

As shown in Figure 3, *Proteobacteria* (22.64% and 31.10%), *Actinobacteria* (20.94% and 14.58%), *Firmicutes* (16.32% and 11.82%), *Chloroflexi* (15.34% and 19.03%), *Bacteroidetes* (12.50% and 14.81%), TM7 (3.66% and 1.76%), *Planctomycetes* (1.91% and 1.42%), and *Euryarchaeota* (1.00% and 1.33%) were dominated in total prokaryotic community and active prokaryotic community in CS. Among them, *Proteobacteria, Actinobacteria, Firmicutes, Chloroflexi*, and *Bacteroidetes* were highly abundant in two prokaryotic communities. Meanwhile, abundances of *Proteobacteria, Chloroflexi, Bacteroidetes*, and *Euryarchaeota* in active prokaryotic community were higher than these in total prokaryotic community. However, abundances of other phyla in active prokaryotic community were lower than these in total prokaryotic community. As shown in Figure 4, *Chitinophagaceae*(family) (4.45%), DRC31(order) (3.99%), and *Comamonadaceae*(family) (3.76%) were the top 3 genera in CS’s total prokaryotic community, and *Allochromatium* (6.61%), *Comamonadaceae*(family) (5.77%) and *Chitinophagaceae*(family) (4.13%) were the top 3 genera in CS’s active prokaryotic community.

As shown in Figure 3, *Chloroflexi* (23.88% and 20.83%), *Bacteroidetes* (20.10% and 17.08%), *Proteobacteria* (13.02% and 22.09%), *Firmicutes* (10.14% and 13.62%), *Verrucomicrobia* (9.21% and 6.73%), *Actinobacteria* (7.98% and 9.16%), WWE1 (4.78% and 3.90%), *Euryarchaeota* (2.90% and 0.43%), *Synergistetes* (1.83% and 0.72%), TM7 (1.18% and 0.94%), and *Planctomycetes* (0.75% and 1.15%) were dominated in total prokaryotic community and active prokaryotic community in DS. Among them, *Chloroflexi, Bacteroidetes, Proteobacteria, Firmicutes, Verrucomicrobia* and *Actinobacteria* were highly abundant in two prokaryotic communities. Meanwhile, abundances of *Proteobacteria, Firmicutes, Actinobacteria*, and *Planctomycetes* in active prokaryotic community were higher than these in total prokaryotic community. However, abundances of other phyla in active prokaryotic community were lower than these in total prokaryotic community. As shown in Figure 4, *Bacteroidales*(order) (6.61%), T78 (5.77%), and LD1-PB3(order) (4.13%) were the top 3 genera in DS’s total prokaryotic community, and *Clostridiaceae*(family) (12.20%), WCHB1-05 (8.82%), and *Bacteroidales*(order) (7.24%) were the top 3 genera in CS’s active prokaryotic community.

## 4. Discussion

The molecular analysis is now central to studies examining the diversity of microorganisms in the environment. Much of what is currently known about the microbial ecology of environment samples (such as soil, water, and sludge) has been inferred from studies targeting DNA. Even though there were studies on analysis of total and active bacterial communities in soil (Baldrian et al., 2012b; Griffiths et al., 2000), seawater (Brettar et al., 2012b), even activated sludge (Gómez-Silván et al., 2014; Maza-Márquez et al., 2016). However, this study detailed analyze the similarities between total and active prokaryotic community in different types sludge samples to evaluate whether understanding the structure-function of microbial community of different types sludge is suitable using DNA-based molecular methods. The current discussion about the prokaryotic community structure of different types sludge based on RNA and DNA is focusing on two key questions: (i) Are there high similarity between total and active prokaryotic communities in different types sludge, and (ii) which type sludge has the highest similarity in all types sludge samples. On top of the two questions remain the detailed understanding of the overall community structure. For this overall structure, the understanding of the core community with the core taxa performing all major functions is essential (Hofle et al., 2008).

### 4.1 The high similarity between total and active prokaryotic communities in sludge

A major goal of this study was to test the similarity between total and active prokaryotic communities in different types sludge samples. The Jarccard’s coefficients of all sludge samples showed that the similarities between total and active procaryotic communities of them were considerable high in this present study. For example, *Firmicutes* and *Proteobacteria* were dominant phyla in PS’s total and active procaryotic communities. A previous study of analysis of bacterial core communities in the central Baltic by comparative RNA–DNA-based fingerprinting showed that large discrepancy between the different types (RNA-, DNA-based fingerprints) of fingerprints in surface water, whereas an unexpected high similarity of both types in the anoxic deep water due to a stable and active core community in this layer (Brettar et al., 2012b). This previous study revealed that low oxidative stress could be a conceivable mechanism explaining the increase of similarity between the RNA-and DNA-based community fingerprints in the anoxic zone (Brettar et al., 2012b). However, in this present study, low oxidative stress was not the conceivable mechanism. In sludge samples, although some of the present bacteria are not as active as others due to the high dynamics, the death rate of most procaryotes may be similar, which explained that the similarity was considerable high in the aerobic and anaerobic sludge samples. This was different from results in soil and surface seawater (Baldrian et al., 2012b; Brettar et al., 2012b). It may be because that sludge is controlled environmental sample, but soil and surface seawater are natural environment sample.

In this present study, procaryotic communities in different types sludge samples were very different, because different type sludge was in different environment and played different roles. For example, *Actinobacteria, Proteobacteria*, and *Chloroflexi* were dominant phyla in excess sludge, because excess sludge was similar to activated sludge. Previous studies have shown that those 3 phyla were dominant in activated sludge from wastewater treatment plants (Gao et al., 2016; Ju et al., 2014; Ju and Zhang, 2015). However, *Verrucomicrobia* and WWE1 were dominant phyla in digested sludge, which were showed in anaerobic reactor conditions (Limam et al., 2014; Riviere et al., 2009).

Moreover, the predominant phylum and genus in total and active procaryotic communities were same in each sludge, and their abundances in active procaryotic community were higher than those in total procaryotic community in all different types sludge samples. It may be because that the predominant phylum and genus are functional microorganism in sludge samples, which is easy to detect using RNA-based illumine MiSeq.

### 4.2 The similarity between total and active prokaryotic communities in running sludge higher than that in idle sludge

More intriguing in our study was that the similarity between the RNA and DNA-based procaryotic communities in running sludge higher than that in idle sludge due that the Jarccard’s coefficient of digested sludge was the highest in all types sludge samples. It might be because that the running sludge (such as digested sludge and ES) played roles in the processes of wastewater treatment and sludge anaerobic digestion. Thus, abundant functional microorganisms existed in the running sludge. For example, Candidatus *M. parvicella*, which might have been responsible for phosphorus removal during sludge bulking in plants with similar configurations (Wang et al., 2014a; Wang et al., 2014b), and *Thauera*, which is affiliated with or closely related to denitrifying bacteria (Wang et al., 2014a), in ES’s active procaryotic community were more abundant than these in ES’s total procaryotic community. An important phenomenon was that abundance of *Methanosaeta*, in digested sludge’s active procaryotic community was 3 times higher than that in digested sludge’s total procaryotic community. However, in the idle sludge, the mortality most of procaryotes greatly increased due to some reasons such as the lack of nutrients.

## 5. Conclusion

Our study demonstrated the high similarity between active and total procaryotic communities in different types sludge samples using RNA-and DNA-based Illumina MiSeq. Meanwhile, the similarity between active and total procaryotic communities in running sludge higher than that in idle sludge. This study showed that the popular DNA-based molecular technology could be applicable to profile functional procaryotic community of sludge samples to a certain degree.

## Acknowledgements

This work was supported by the National Key R&D Program of China(2018YFC0407803), the Natural Science Research Project of Henan Educational Committee (18B610010), the Talent-Recruiting Program of Zhoukou Normal University (ZKNUC2017037), and the Yellow River Institute of Hydraulic Research (HKY-jyxm-2017-04).

## References

Baldrian, P, Kolařík, M, Stursová, M, Kopecký, J, Valášková, V, Větrovský, T, Zifčáková, L, Snajdr, J, Rídl, J, Vlček, C, 2012a. Active and total microbial communities in forest soil are largely different and highly stratified during decomposition. The ISME Journal 6:248–258.

Baldrian, P, Kolařík, M, Štursová, M, Kopecký, J, Valášková, V, Větrovský, T, Žifčáková, L, Šnajdr, J, Rídl, J, Vlček, Č, 2012b. Active and total microbial communities in forest soil are largely different and highly stratified during decomposition. The ISME Journal 6:248–258.

Braguglia, C, Gagliano, M, Rossetti, S, 2012. High frequency ultrasound pretreatment for sludge anaerobic digestion: effect on floc structure and microbial population. Bioresource Technology 110:43–49.

Brettar, I, Christen, R, Höfle, M G, 2012a. Analysis of bacterial core communities in the central Baltic by comparative RNA-DNA-based fingerprinting provides links to structure-function relationships. The ISME Journal 6:195–212.

Brettar, I, Christen, R, Höfle, M G, 2012b. Analysis of bacterial core communities in the central Baltic by comparative RNA–DNA-based fingerprinting provides links to structure–function relationships. The ISME Journal 6:195–212.

Cui, R, Jahng, D, 2006. Enhanced methane production from anaerobic digestion of disintegrated and deproteinized excess sludge. Biotechnology Letters 28:531–538.

Edgar, R C, Haas, B J, Clemente, J C, Quince, C, Knight, R, 2011. UCHIME improves sensitivity and speed of chimera detection. Bioinformatics 27:2194–2200.

Erhart, R, Bradford, D, Seviour, R J, Amann, R, Blackall, L L, 1997. Development and use of fluorescent in situ hybridization probes for the detection and identification of “Microthrix parvicella” in activated sludge. Systematic and Applied Microbiology 20:310–318.

Franchi, O, P Bovio, E Ortega-Martínez, F Rosenkranz, and R Chamy. 2018. Active and total microbial community dynamics and the role of functional genes bamA and mcrA during anaerobic digestion of phenol and p-cresol. Bioresource Technology 264: 290–298.

Gómez-Silván, C, Arévalo, J, González-López, J, Rodelas, B, 2014. Exploring the links between population dynamics of total and active bacteria and the variables influencing a full-scale membrane bioreactor (MBR). Bioresource Technology 162:103–114.

Gao, P, Xu, W, Sontag, P, Li, X, Xue, G, Liu, T, Sun, W, 2016. Correlating microbial community compositions with environmental factors in activated sludge from four full-scale municipal wastewater treatment plants in Shanghai, China. Applied Microbiology and Biotechnology 100:1–11.

Griffiths, R I, Whiteley, A S, O’Donnell, A G, Bailey, M J, 2000. Rapid method for coextraction of DNA and RNA from natural environments for analysis of ribosomal DNA-and rRNA-based microbial community composition. Applied and environmental microbiology 66:5488–5491.

Hofle, M, Kirchman, D L, Christen, R, Brettar, I, 2008. Molecular diversity of bacterioplankton: link to a predictive biogeochemistry of pelagic ecosystem. Aquatic Microbial Ecology 53:39–48.

Ju, F, Guo, F, Ye, L, Xia, Y, Zhang, T, 2014. Metagenomic analysis on seasonal microbial variations of activated sludge from a full-scale wastewater treatment plant over 4 years. Environmental Microbiology Reports 6:80–89.

Ju, F, Zhang, T, 2015. Bacterial assembly and temporal dynamics in activated sludge of a full-scale municipal wastewater treatment plant. The ISME Journal 9:683–695.

Kim, J, Park, C, Kim, T H, Lee, M, Kim, S, Kim, S W, Lee, J, 2003. Effects of Various Pretreatments for Enhanced Anaerobic Digestion with Waste Activated Sludge. Journal of Bioscience & Bioengineering 95:271–275.

Korzeniewska, E., and M. Harnisz. 2018. Relationship between modification of activated sludge wastewater treatment and changes in antibiotic resistance of bacteria Science of the Total Environment 639:304–315.

Lanzén, A., Jørgensen, S. L., Bengtsson, M. M., Jonassen, I., Øvreås, L., & Urich, T., 2011. Exploring the composition and diversity of microbial communities at the Jan Mayen hydrothermal vent field using RNA and DNA. FEMS microbiology ecology 77:577–589.

Limam, R D, Chouari, R, Mazéas, L, Wu, T D, Li, T, Grossin-Debattista, J, Guerquin-Kern, J L, Saidi, M, Landoulsi, A, Sghir, A, 2014. Members of the uncultured bacterial candidate division WWE1 are implicated in anaerobic digestion of cellulose. MicrobiologyOpen 3:157–167.

Liu, W T, Marsh, T L, Cheng, H, Forney, L J, 1997. Characterization of microbial diversity by determining terminal restriction fragment length polymorphisms of genes encoding 16S rRNA. Applied and Environmental Microbiology 63:4516–4522.

Lozupone, C, Lladser, M E, Knights, D, Stombaugh, J, Knight, R, 2011. UniFrac: An effective distance metric for microbial community comparison. The ISME Journal 5:169–172.

Maza-Márquez, P, Vílchez-Vargas, R, Boon, N, González-López, J, Martínez-Toledo, M, Rodelas, B, 2016. The ratio of metabolically active versus total Mycolata populations triggers foaming in a membrane bioreactor. Water Research 92:208–217.

Mikkonen, A, Santalahti, M, Lappi, K, Pulkkinen, A M, Montonen, L, Suominen, L, 2014. Bacterial and archaeal communities in long-term contaminated surface and subsurface soil evaluated through coextracted RNA and DNA. FEMS Microbiology Ecology 90:103–114.

Mossakowska, A, Hellström, B G, Hultman, B, 1998. Strategies for sludge handling in the Stockholm region. Water Science & Technology 38:111–118.

Neilson, A, 1978. The occurrence of aeromonads in activated sludge: isolation of Aeromonas sobria and its possible confusion with Escherichia coli. Journal of Applied Bacteriology 44:259–264.

Rincón, B, Borja, R, González, J, Portillo, M, Sáiz-Jiménez, C, 2008. Influence of organic loading rate and hydraulic retention time on the performance, stability and microbial communities of one-stage anaerobic digestion of two-phase olive mill solid residue. Biochemical Engineering Journal 40:253– 261.

Riviere, D, Desvignes, V, Pelletier, E, Chaussonnerie, S, Guermazi, S, Weissenbach, J, Li, T, Camacho, P, Sghir, A, 2009. Towards the definition of a core of microorganisms involved in anaerobic digestion of sludge. The ISME Journal 3:700–714.

Rulkens, W, 2007. Sewage Sludge as a Biomass Resource for the Production of Energy: Overview and Assessment of the Various Options†. Energy & Fuels 22:9–15.

Schloss, P D, Westcott, S L, Ryabin, T, Hall, J R, Hartmann, M, Hollister, E B, Lesniewski, R A, Oakley, B B, Parks, D H, Robinson, C J, 2009. Introducing mothur: open-source, platform-independent, community-supported software for describing and comparing microbial communities. Applied and Environmental Microbiology 75:7537–7541.

Schuppler, M, Mertens, F, Schön, G, Göbel, U B, 1995. Molecular characterization of nocardioform actinomycetes in activated sludge by 16S rRNA analysis. Microbiology 141:513–521.

Sudhagar, Arun, Gokhlesh Kumar, and Mansour El-Matbouli. 2018. Transcriptome Analysis Based on RNA-Seq in Understanding Pathogenic Mechanisms of Diseases and the Immune System of Fish: A Comprehensive Review. International Journal of Molecular Sciences 19: 245–251.

Sundberg, C, Al-Soud, W A, Larsson, M, Alm, E, Yekta, S S, Svensson, B H, Sørensen, S J, Karlsson, A, 2013. 454-Pyrosequencing Analyses of Bacterial And Archaeal Richness In 21 Full-Scale Biogas Digesters. FEMS Microbiology Ecology 85:612–626.

Supaphol, S, Jenkins, S N, Intomo, P, Waite, I S, O’Donnell, A G, 2011. Microbial community dynamics in mesophilic anaerobic co-digestion of mixed waste. Bioresource Technology 102:4021–4027.

Vanwonterghem, I, Jensen, P D, Dennis, P G, Hugenholtz, P, Rabaey, K, Tyson, G W, 2014. Deterministic processes guide long-term synchronised population dynamics in replicate anaerobic digesters. The ISME Journal 8:2015–2028.

Wagner, M, Loy, A, Nogueira, R, Purkhold, U, Lee, N, Daims, H, 2002. Microbial community composition and function in wastewater treatment plants. Antonie Van Leeuwenhoek 81:665–680.

Wang, J, Li, Q, Qi, R, Tandoi, V, Yang, M, 2014a. Sludge bulking impact on relevant bacterial populations in a full-scale municipal wastewater treatment plant. Process Biochemistry 49:2258–2265.

Wang, J, Qi, R, Liu, M, Li, Q, Bao, H, Li, Y, Wang, S, Tandoi, V, Yang, M, 2014b. The potential role of Candidatus ‘Microthrix parvicella’ in phosphorus removal during sludge bulking in two full-scale enhanced biological phosphorus removal plants. Water Science and Technology 70:367–375.

Wang, Y, Sho, M, Naoto, O, Takeshi, F, 2011. A survey of the cellular responses in Pseudomonas putida KT2440 growing in sterilized soil by microarray analysis. FEMS Microbiology Ecology 78:220–232.

Wang, Weifeng, Yu Jie, Cui Yang, Jun He, Xue Peng, Cao Wan, Hongmei Ying, Wenkang Gao, Yingchao Yan, and Hu Bo. 2018. Characteristics of fine particulate matter and its sources in an industrialized coastal city, Ningbo, Yangtze River Delta, China. Atmospheric Research 203:105–117.

Weemaes, M, Grootaerd, H, Simoens, F, Verstraete, W, 2000. Anaerobic digestion of ozonized biosolids. Water Research 34:2330–2336.

Wong, M T, Zhang, D, Li, J, Hui, R K H, Tun, H M, Brar, M S, Park, T J, Chen, Y, Leung, F C, 2013. Towards a metagenomic understanding on enhanced biomethane production from waste activated sludge after pH 10 pretreatment. Biotechnology and Biofuels 6:38–45.

Ye, L, Zhang, T, 2011. Pathogenic bacteria in sewage treatment plants as revealed by 454 pyrosequencing. Environmental Science & Technology 45:7173–7179.

Ye, L, Zhang, T, 2013. Bacterial communities in different sections of a municipal wastewater treatment plant revealed by 16S rDNA 454 pyrosequencing. Applied Microbiology and Biotechnology 97:2681– 2690.

Zakrzewski, M, Goesmann, A, Jaenicke, S, Jünemann, S, Eikmeyer, F, Szczepanowski, R, Al-Soud, W A, Sørensen, S, Pühler, A, Schlüter, A, 2012. Profiling of the metabolically active community from a production-scale biogas plant by means of high-throughput metatranscriptome sequencing. Journal of Biotechnology 158:248–258.

Zhang, D, Chen, Y, Zhao, Y, Zhu, X, 2010. New sludge pretreatment method to improve methane production in waste activated sludge digestion. Environmental Science & Technology 44:4802–4808.

Zhang, T, Shao, M-F, Ye, L, 2012. 454 Pyrosequencing reveals bacterial diversity of activated sludge from 14 sewage treatment plants. The ISME Journal 6:1137–1147.

